# BMSCs differentiated into neurons, astrocytes and oligodendrocytes alleviated the inflammation and demyelination of EAE mice models

**DOI:** 10.1101/2020.11.16.384354

**Authors:** Guo-yi Liu, Yan Wu, Fan-yi Kong, Shu Ma, Li-yan Fu, Jia Geng

## Abstract

Multiple sclerosis (MS) is a complex, progressive neuroinflammatory disease associated with autoimmunity. Currently, effective therapeutic strategy was poorly found in MS. Experimental autoimmune encephalomyelitis (EAE) is widely used to study the pathogenesis of MS. Previous studies have shown that bone marrow mesenchymal stem Cells (BMSCs) transplantation could treat EAE animal models, but the mechanism was divergent. Here, we systematically evaluated whether BMSCs can differentiate into neurons, astrocytes and oligodendrocytes to alleviate the symptoms of EAE mice. We used Immunofluorescence staining to detect MAP-2 neurons marker, GFAP astrocytes marker, and MBP oligodendrocytes marker expression to evaluate whether BMSCs can differentiate. The effect of BMSCs transplantation on inflammatory cell invasion and demyelination in EAE mice were detected by Hematoxylin-Eosin (H&E) and Luxol Fast Blue (LFB) staining. Inflammatory factors expression was detected by ELISA and RT-qPCR. Our results showed that BMSCs could be induced to differentiate into neuron cells, astrocytes and oligodendrocyte in vivo and in vitro. In addition, BMSCs transplant improved the survival rate and weight, and reduced neurological function scores and disease incidence of EAE mice. Moreover, BMSCs transplant alleviated the inflammation and demyelination of EAE mice. Finally, we found that BMSCs transplantation down-regulated the expression levels of pro-inflammatory factors TNF-α, IL-1β and IFN-γ, and up-regulated the expression levels of anti-inflammatory factors IL-10 and TGF-β. In conclusion, this study found that BMSCs could alleviate the inflammatory response and demyelination in EAE mice, which may be achieved by the differentiation of BMSCs into neurons, astrocytes and oligodendrocytes in EAE mice.

## 1. Introduction

Multiple sclerosis (MS) is a chronic neuroinflammatory disease that is associated with autoimmunity in central nervous system (CNS) ^[1, 2]^. Generally speaking, it is characterized by axon damage, demyelination, inflammatory infiltration and progressive neurological damage, eventually leading to disability ^[3]^. Moreover, MS is thought to be a multifocal demyelination disease in CNS ^[4]^. Currently, MS mainly depends on three types of drug treatment: disease-modifying drugs (DMD) specifically designed for MS, corticosteroids for acute exacerbations, and drugs for symptomatic control. However, it is less evident that drugs are effective in the progression of MS ^[5]^. Thus, to find an effective therapeutic strategy is very important for MS in clinical. Experimental autoimmune encephalomyelitis (EAE) is widely used to study the pathogenesis of MS, which represents both pathological and features of MS ^[6]^. To some extent, EAE could effectively elucidate various pathological processes in MS, such as inflammation demyelination, axonal lesions ^[7]^.Bone marrow mesenchymal stem Cells (BMSCs) are non-hematopoietic stromal cells that derived from bone marrow^[8]^. BMSCs differentiate into various cell types, which contribute to regeneration of tissues^[9–11]^. BMSCs also play immunomodulatory role through inhibiting T-cell activities^[12]^. In addition, BMSCs secrete growth factors to regulate hematopoietic stem/progenitor cell proliferation and differentiation^[13]^. Recently, some researchers have shifted the focus to stem cell-based therapy in many diseases, including nervous system disease, indicating that stem cell may be potentially amenable to therapeutic manipulation for clinical application in MS^[14, 15]^. In the acute and subacute phases, the tissues were selectively target damaged by intravenous injection of BMSCs, which improved the recovery of neurological function, decreased inflammatory response and demyelination after EAE^[16]^ However, whether BMSCs have a therapeutic effect on EAE mice through differentiation remains to be studied. In addition, it is now recognized that the interaction of neurons, astrocytes and oligodendrocytes plays an important regulatory role in remyelination and the development of and MS[17, 18]. Therefore, we explored whether the transplanted BMSCs can differentiate into neurons, astrocytes and oligodendrocytes to alleviate the development of MS. In our study, we found that BMSCs could differentiate into neurons, astrocytes and oligodendrocytes. At the same time, we established the EAE animal model by injected subcutaneously with MOG35-55 peptide. In addition, we performed BMSCs transplant and determined the effects of BMSCs on the progression of EAE in mice.

## 2. Materials and Methods

### 2.1 Animal

C57BL/6 mice (n=30) were purchased from Kunming Medical University. The mice were housed in sterile, constant temperature rooms at Kunming Medical University with a 12h/12h light/dark cycle, with free access to food and water. All animal experiments were conducted according to the ARRIVE guidelines, and were performed in accordance with Ethics Committee of Kunming Medical University and approved by Ethics Committee of Kunming Medical University. All measures had been taken to minimize the suffering of animals. Compared to the starting point of EAE immunization, no animal lost more than 20% of its body weight and no neurological score exceeded 4. The health of the mice was monitored at least once a day, and no accidental deaths were observed. Euthanasia was carried out in a CO_2_ chamber, gradually filled with CO_2_ before 4 of mice neurological score, and then bled. All animal facility staff and researchers had followed required courses and obtained certificates to conduct animal research.

### 2.2 EAE model

EAE model was induced by 125μg of MOG35-55 peptide diluted with PBS(PH=7.2) to 0.01mol/mL, and mixed with complete Freund’s adjuvant (CFA; Shanghai, China) including Mycobacterium tuberculosis. Each mouse was injected subcutaneously at one point in the groin and three points in the back. Subsequently, the mice were injected with 200ng pertussis toxin (Thermo Fisher Scientific;Beijing, China). After 48h, the mice were injected with 200 ng pertussis toxin again. On the 7th day, we also injected with 200 ng pertussis toxin into the mice. 30 mice were randomly divided into two groups. BMSCs were injected into the lateral ventricle (Start with the dura mater; before and after the halogen: 0.6mnl; opening: 1.5mm; depth: 1.7mm) of EAE mice by brain stereotactic technology.

### 2.3 Isolation and identification of primary BMSCs

According to the method of Gao et al[19]., BMSCs were isolated from male mice anesthetized (1% sodium pentobarbital, 40 mg/kg intraperitoneal injection). BMSCs were cultured at 37°C and 5% CO_2_ with DMEM/F12 medium (Gibco, USA) supplemented with 10% fetal bovine serum (FBS; Gibco, USA),100 U/mL penicillin and 100 U/mL streptomycin after the culture medium is washed. In our previous study, we have evaluated phenotype of BMSCs by flow cytometric analysis.

### 2.4 Induced BMSCs to differentiate into neurons, astrocytes and oligodendrocytes

BMSCs from the fifth generation were seeded in 6-well plates coated with poly ornithine and laminin at a density of 2×10^5^ cells/mL. After 24 h, respectively, the DMEM/F12 medium were replaced with complete culture medium (Gibco, USA) containing 10 ng/mlL bFGF to induced as neurons, or 1% N_2_ and 2% B27 to induced as astrocytes, or 2 mM glutamine, 50 U/mL penicillin, 50 U/mL streptomycin, and 60 μg/mL T3 to induced as oligodendrocytes. The complete culture medium was changed every 2 to 3 days. BMSCs in control group were not induced.

### 2.5 Clinical signs measurement

During the experiment, the weight, neurobehavioral score, disease incidence and survival time of the mice were monitored by two independent observers in a blind manner until the 36th day after immunization. The neurobehavioral score is defined by the following scale: no clinical symptoms=0; wobbled gait or loss of tail tension=1; tail weakness, hind limb weakness=2; hind limb paralysis=3; hind limb paralysis, forelimb paralysis or Forelimb weakness with bowel dysfunction=4; dying or death=5; the symbol between them is ±0.5[20]. In our preliminary experiments, the peak of the disease is from the 9th to the 15th day after immunization. After 16th days after immunization, the mice were euthanized by cervical dislocation. We collected serum from the mice and took cortex and hippocampus of brain tissues and lumbar spinal cord for the following experiments.

### 2.6 Immunofluorescence staining (IF)

After fixed with 4% paraformaldehyde, the cells/brain sections were penetrated with PBS containing 0.4% Triton X-100. Then, the cells/brain sections were blocked with 5% Bovine Serum Albumin (BSA; Beijing; China) at room temperature for 1h. The primary antibodies were incubated overnight at 4℃. The secondary antibodies with fluorophores (1:1000, KPL) were incubated at 4℃ for 1h. Then, nuclei staining with DAPI followed by capturing using a microscope (Olympus, Japan). We calculated the ratio of BMSCs to differentiate into neurons, astrocytes and oligodendrocytes, respectively. The primary antibodies were shown in Table.1.

**Table.1.** Primary antibodies.

| Primary antibodies | Company | Dilution |
| --- | --- | --- |
| MAP-2 | Abcam | 1:600 |
| βIII-tubulin | Abcam | 1:400 |
| GFAP | Abcam | 1:300 |
| MBP (Myelin Basic Protein) | Abcam | 1:400 |
| O4(Oligodendrocyte Marker) | Sigma | 1:500 |

### 2.7 Luxol Fast Blue staining (LFB)

Spinal cord sections of Lumbar spine were placed in a xylene solution for gradient hydration. Then, the sections were put into Luxol fast blue dye solution (Sigma, USA) in a 50-65℃ incubator overnight. Stained sections were taken out and passed through alcohol, water, and then added the color separation solution. Sections were washed 3 times with water after differentiation, and then counterstained or gradient dehydrated., Each mice analyzed 3 histological sections and calculated the average score in our experiment. Standard score for the degree of demyelination: none =0; rare lesions=1; demyelination of a few areas=2; demyelination of large areas or fusion areas=3[21].

### 2.8 Hematoxylin-Eosin staining (H&E)

The mice were sacrificed and perfused with normal saline and 4% paraformaldehyde through the blood vessel. The lumbar spinal cord was fixed with 4% paraformaldehyde for 24 hours, embedded in paraffin, and cut into 4-5 μm thick sections. H&E staining was performed according to the manufacturer’s instructions to assess the degree of inflammatory cell infiltration. In our experiment, each mice analyzed 3 histological sections and calculated the average score. Standard scores for the degree of inflammatory infiltration: no infiltrating cells=0; a small amount of scattered infiltrating cells=1; inflammatory infiltrating tissue around blood vessels=2; extensive perivascular scar infiltration=3[21].

### 2.9 RNA extraction and Real-time quantitative PCR (RT-PCR)

According to the instructions, total RNA of cells and brain tissues were extracted using Trizol Reagent (Lifetech, USA). We followed the instructions of the FastKing RT Kit (Fermentas; Shanghai, China) to synthesize the first strand of cDNA. Then, we performed quantitative PCR by SYBR Green master mix (KAPA; Shanghai, China). The primers were designed using beacon designer 7.90. The primer sequences were listed in Table.2. All experimental results were analyzed by 2^−△△Ct^ method.

**Table.2.**
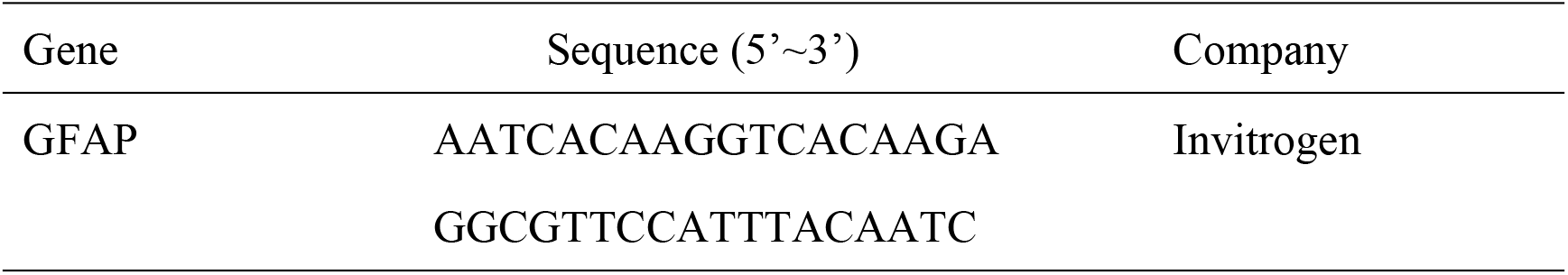
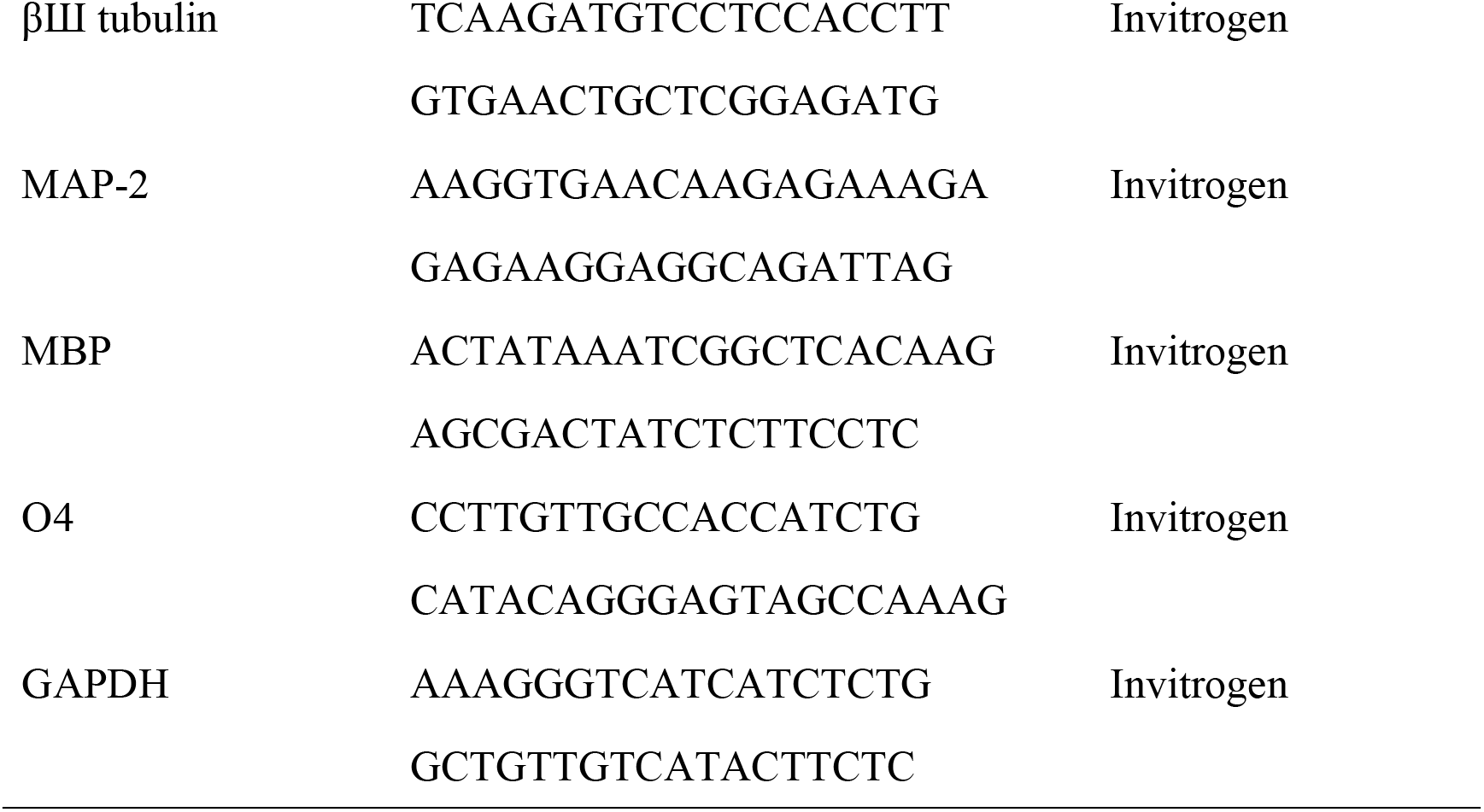
Primer sequences

### 2.10 ELISA Assay

The levels of inflammatory factors TNF-α, IL-1β and IFN-γ, and anti-inflammatory factors IL-10 and TGF-β were detected by ELISA. According to the instructions of ELISA Kit, the supernatant was collected by centrifugation homogenate (5000rpm for 15 minutes at 4 ℃). The levels TNF-α, IL-1β, IFN-γ, IL-10 and TGF-β were determined by ELISA Kit. We constructed standard curves by standard samples. Quantification of ELISA results were performed at 450nm using an ELISA plate reader (Spectra Max 190, Molecular Devices, USA).

### 2.11 Statistical analysis

GraphPad Prism 7 software (GraphPad, USA) was used to conduct statistical analysis. One-way ANOVA and t test were used to analyze data. Data were presented as mean ± standard deviation (SD) and P value<0.05 were considered as significant results.

## 3. Results

### 3.1 Identification of primary BMSCs

Before the BMSCs experiment, we used an inverted phase contrast microscope and flow cytometry to identify the morphology and marker expression levels of BMSCs. Observed under an inverted phase contrast microscope, the BMSCs cells of the P1 generation adhered to the wall and presented a polygonal shape. After growing to the P3 generation, the overall BMSCs cells showed a swirl shape at high density, while BMSCs cells gradually showed a fibroblast-like spindle shape at low density (Fig. 1A). The above observation results are consistent with the basic morphological characteristics of BMSCs. Furthermore, we detected the levels of BMSCs related markers CD29, CD90, CD34 and CD45 by flow cytometry. The result showed that cultured BMSCs expressed positive CD29^+^ (94.55%) and CD90^+^ (94.67%), and negative CD34^−^ (1.43%) and CD45^−^ (0.83%) on the surface of BMSCs (Fig.1B). All the results were consistent with the previous studies^[22, 23]^.

**Figure 1.**
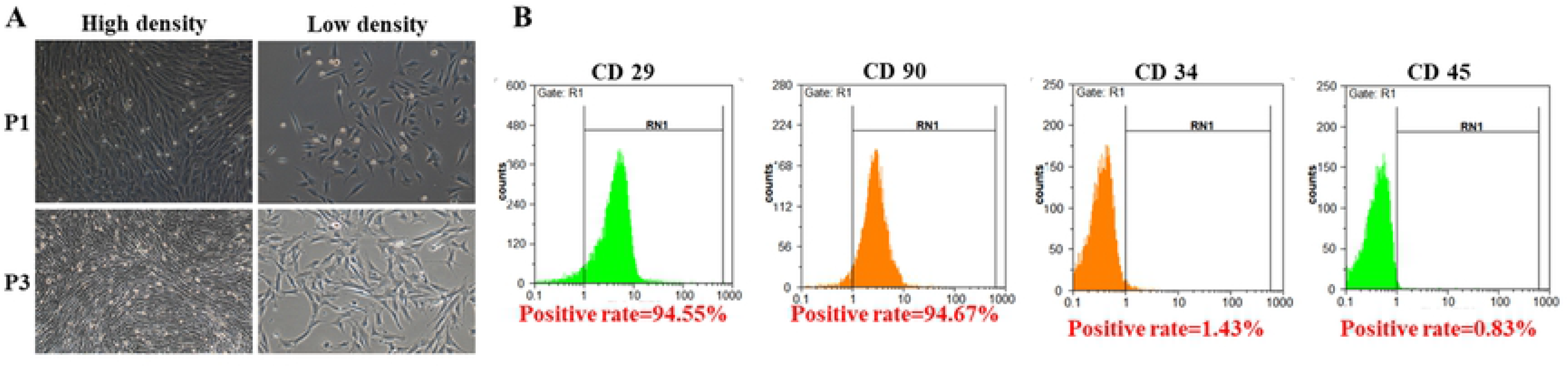
Identification of primary BMSCs. (A)BMSCs morphology was assessed by microscopy (scale bar=100 μm). (B) CD29, CD34, CD45 and CD90 surface marker expression was determined by flow cytometry.

### 3.2 BMSCs were induced to differentiate into neurons, astrocytes and oligodendrocytes

To verify that BMSCs can differentiate into neurons, astrocytes and oligodendrocytes, we induced the culture of BMSCs, and used IF and RT-qPCR to detect the expression level of neurons markers (MAP-2 and βIII-tubulin), astrocytes markers (GFAP) and oligodendrocytes markers (MBP and O4), respectively. The results of IF showed that MAP-2 and βIII-tubulin were expressed in BMSCs cultured in neurons induction medium (Fig. 2A). GFAP was expressed in BMSCs cultured in astrocytes induction medium (Fig. 2C). MBP and O4 were expressed in BMSCs cultured in oligodendrocytes induction medium (Fig. 2E). In addition, the RT-qPCR results showed that compared with the NC group treated with PBS, the expression levels of MAP-2 and βIII-tubulin in the Induced neurons group (Fig. 2B), GFAP in the Induced astrocytes group (Fig. 2D), and MBP and O4 in the Induced neurons group (Fig. 2F) significantly upregulated. It could be seen from the above experimental results that BMSCs could be induced to differentiate into neurons, astrocytes and oligodendrocytes in vitro.

**Figure 2.**
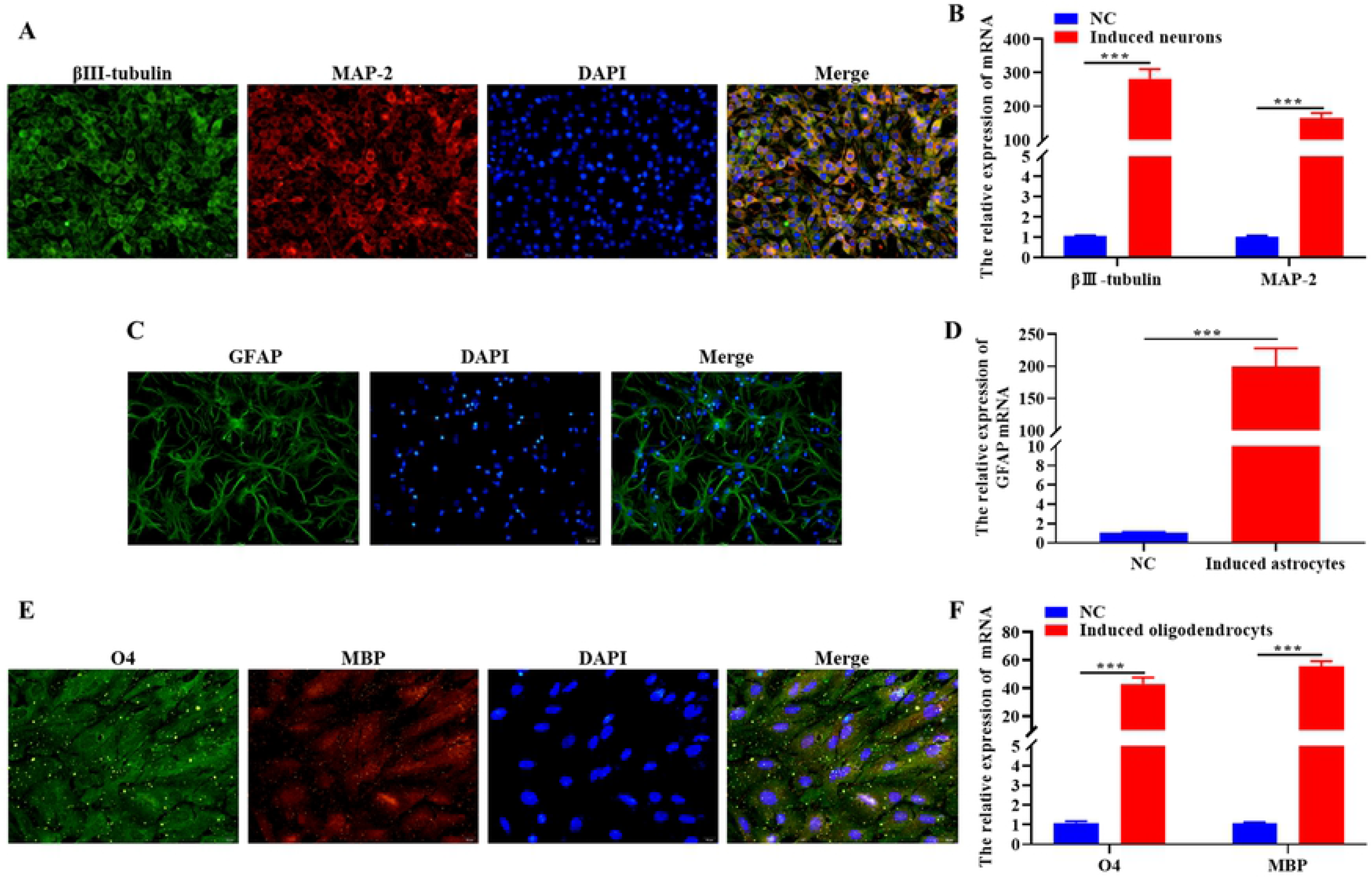
BMSCs were induced to differentiate into neurons, astrocytes and oligodendrocytes. The position and quantification of MAP-2 and βIII-tubulin (A), GFAP (C), MBP and O4 (E) were detected by IF staining. The expression levels of MAP-2 and βIII-tubulin (B), GFAP (D), MBP and O4 (F) mRNA were detected by RT-qPCR. Original magnification: 40×. Data were expressed as mean ± SD. Comparison with NC group, \*\**P*<0.01, \*\*\**P*<0.001.

### 3.3 The transplanted BMSCs differentiated into neurons, astrocytes and oligodendrocytes in the EAE mice

To verify that BMSCs can differentiate into neurons, astrocytes and oligodendrocytes in vivo, we transplanted GFP-labeled BMSCs into EAE mice model by brain stereotactic technology, and observed the differentiation effect of BMSCs in hippocampus and cortex through IF. The results of the IF experiment showed that the GFP fluorescence signal could be observed in the EAE+BMSCs group, while the EAE group had no GFP fluorescence signal at all. In addition, after Image J fusion, it was found that the expression positions of MAP-2, βIII-tubulin, GFAP, MBP and O4 overlap with the fluorescent signal positions of GFP in the hippocampus and cortex (Fig. 3A-3C). The above experiments show that transplanted BMSCs could differentiate into neurons, astrocytes and oligodendrocytes in the EAE mice. Furthermore, we conducted a statistical analysis of the experimental results of IF and found that the fluorescence intensities of MAP-2, βIII-tubulin, GFAP, MBP and O4 in the hippocampus and cortex of the BMSCs transplantation group were significantly higher than those in EAE group (Fig.3D-3F). It is suggested that the transplanted BMSCs could increase the number of neurons, astrocytes and oligodendrocytes in the hippocampus and cortex. This result was at least partly due to the differentiation of BMSCs into neurons, astrocytes and oligodendrocytes.

**Figure 3.**
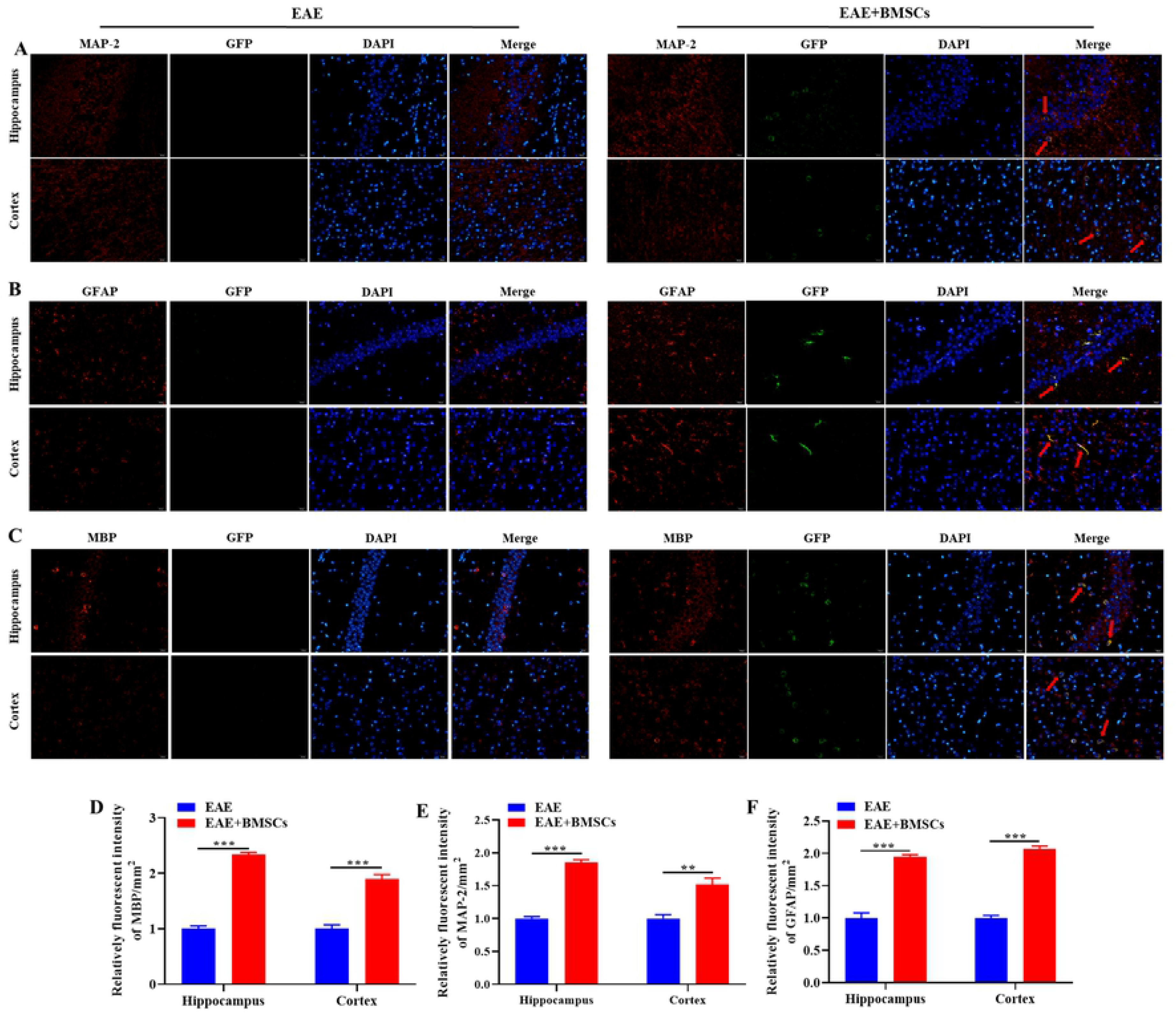
The transplanted BMSCs differentiated into neurons, astrocytes and oligodendrocytes in the EAE mice. The position and quantification of GFP with MAP-2 (A), GFAP (B), MBP (C) were detected by IF staining. The fluorescence intensity of MAP-2 (D), GFAP (E), MBP (F) were detected by Image J software. Original magnification: 40×. Data were expressed as mean ± SD. Comparison with EAE group, \*\**P*<0.01, \*\*\**P*<0.001.

### 3.4 Effect of BMSCs transplantation on clinical signs during EAE progression in mice

To determine whether the differentiation of BMSCs can alleviate the clinical symptoms of EAE, we performed survival analysis, disease incidence analysis, neurobehavioral score and weight measurement on EAE mice and BMSCs transplanted EAE mice. A higher neurobehavioral score indicates worsening of motor dysfunction. Higher disease incidence and lower body weight indicate an increase in disease severity.

The mice survival line chart shown that the survival rate of the mice in the EAE group was significantly lower than that of the mice in the EAE+BMSCs group (Fig. 4A). In addition, the results of the line graph of disease incidence shown that mice in the EAE group began to develop disease on the 6th day after immunization, and by the 15th day, all the mice in the EAE group developed disease (Fig. 4B). However, the mice in the BMSCs treatment group recovered from the disease on the 14th day after immunization and returned to their normal state on the 21st day. The neurobehavioral score of the EAE group increased rapidly after immunization, and reached a peak on the 16th day, and BMSCs treatment could significantly reduce the neurobehavior score of the EAE in mice (Fig. 4C). Furthermore, the mice decreased significantly on the 10th day after immunization in the EAE group. Interestingly, the body weight of mice had been showing an upward trend in the EAE+BMSCs group (Fig. 4D). In summary, the differentiation of transplanted BMSCs could alleviate clinical signs during EAE progression in mice.

**Figure 4.**
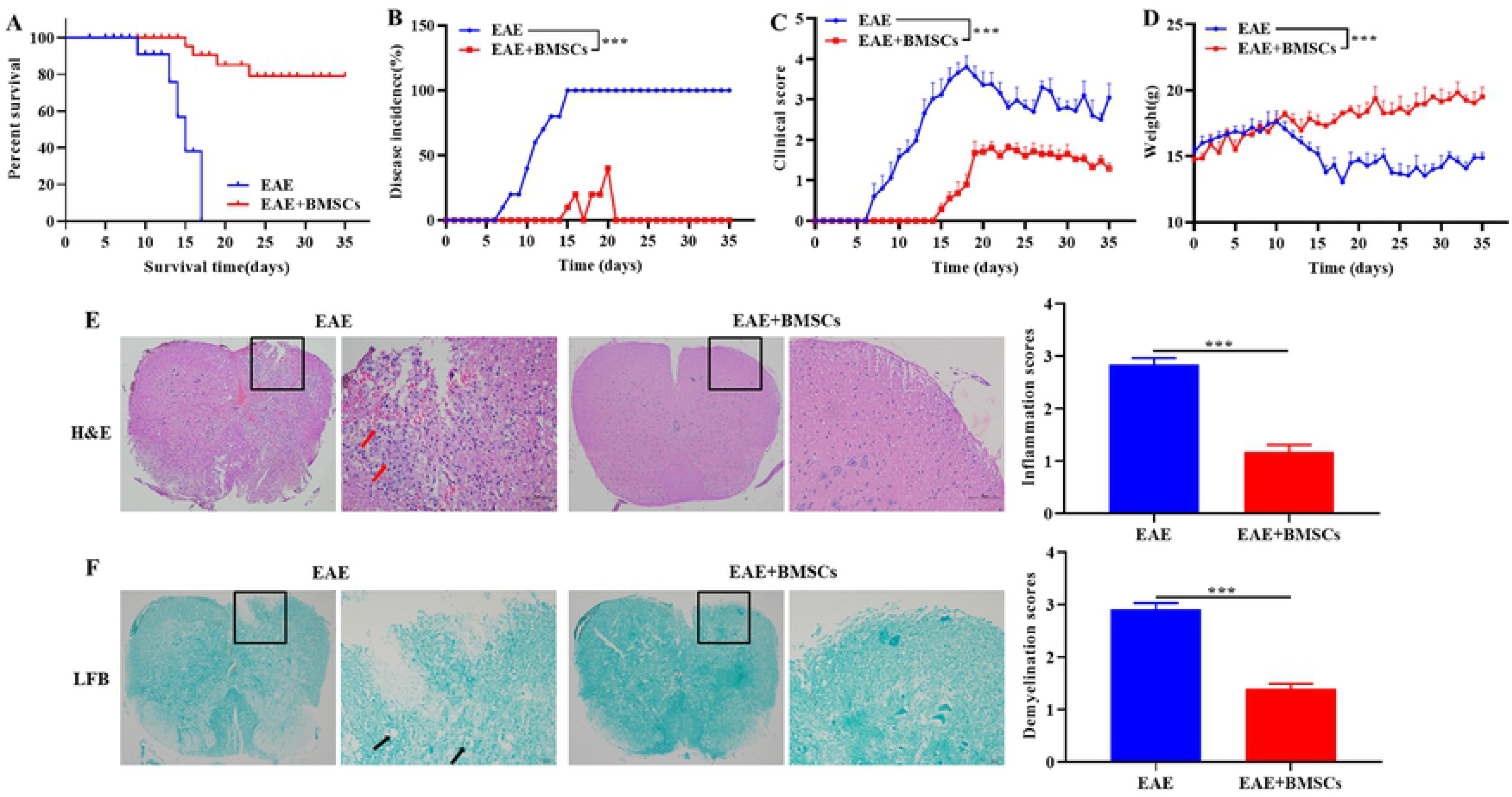
Effect of BMSCs transplantation on clinical signs and histological changes during EAE progression in mice. (A) Survival curve of percent survival of mice in EAE groups and EAE+BMSCs groups. (B) Analysis of disease incidence of mice in EAE groups and EAE+BMSCs groups. (C) Neurobehavioral score of mice in EAE groups and EAE+BMSCs groups. (D) Changes in body weight of mice in EAE groups and EAE+BMSCs groups. (E) H&E staining to observe the infiltration of inflammatory cells in the lumbar spinal cord of mice, and to score and count. (B) LFB staining to observe the demyelination in the lumbar spinal cord of mice, and to score and count. Original magnification: 4× or 20×. Data were expressed as mean ± SD. Comparison with EAE group, \*\*\**P*<0.001.

**Figure 5.**
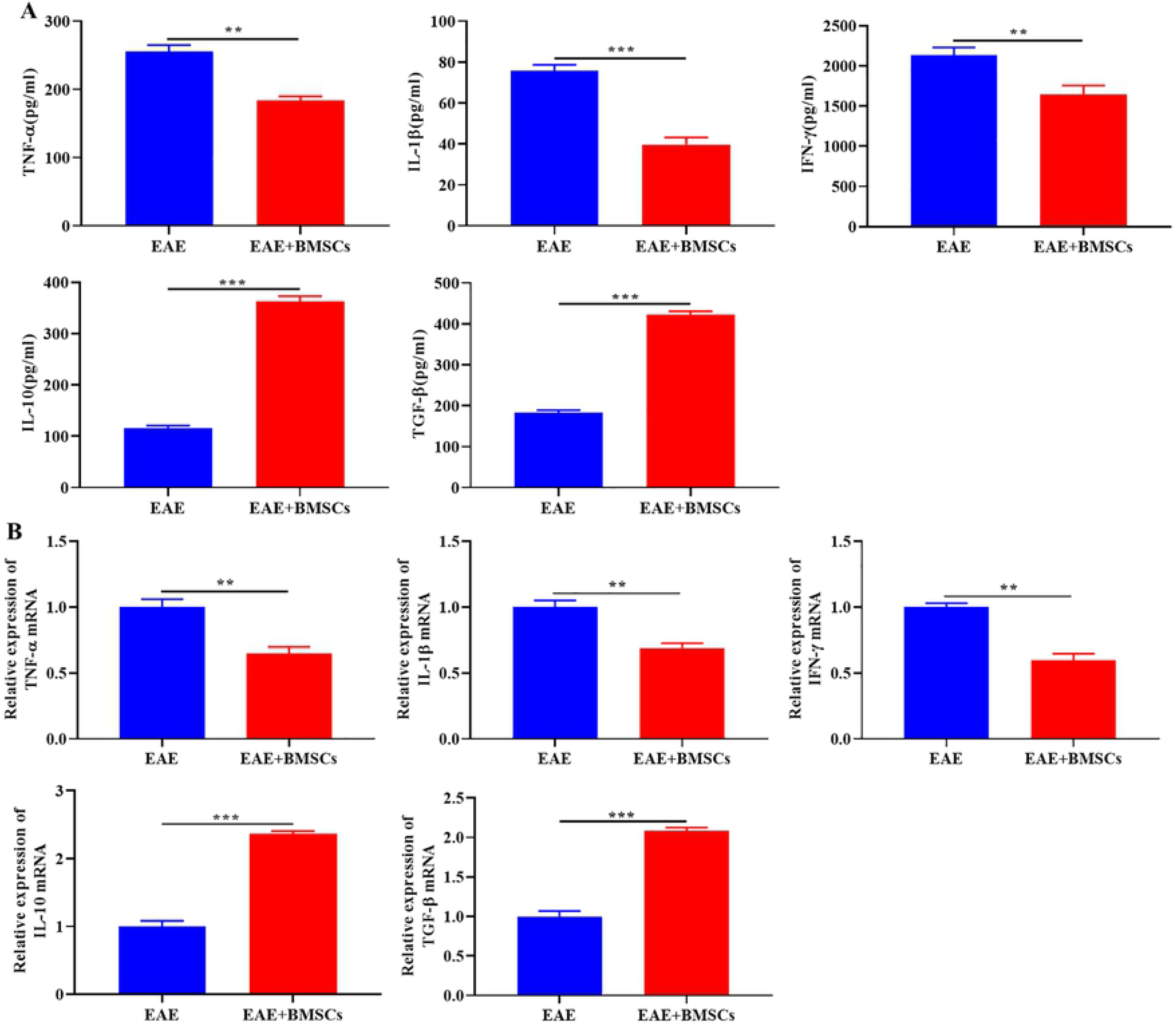
Effect of BMSCs transplantation on inflammatory factor expression during EAE progression in mice. (A) The levels of pro-inflammatory factors TNF-α, IL-1β and IFN-γ and anti-inflammatory factors IL-10 and TGF-β were detected by ELISA. (B) The expression levels of pro-inflammatory factors TNF-α, IL-1β and IFN-γ and anti-inflammatory factors IL-10 and TGF-β mRNA were detected by RT-qPCR. Data were expressed as mean ± SD. Comparison with EAE group, \*\**P*<0.01, \*\*\**P*<0.001.

### 3.5 Effect of BMSCs transplantation on histological changes during EAE progression in mice

We sacrificed the mice on the 16th day after immunization, collected the lumbar spinal cord tissues of mice for histological analysis to analyze the progression of the disease at the level of CNS damage in the EAE group and EAE+BMSCs group. The results of H&E (Fig.4E) and LFB (Fig.4F) shown the inflammatory cell infiltration and demyelination status of mice in the EAE group and EAE+BMSCs group. The results shown that a lot of inflammatory cells gathered around the small blood vessels in the lumbar spinal cord of the EAE group, and the degree of demyelination of the lumbar spinal cord of the EAE group was significantly higher than that of EAE+BMSCs. However, after BMSCs transplantation, the degree of infiltration of inflammatory cells in the lumbar spinal cord of the EAE+BMSCs group was alleviated, and the degree of demyelination was also significantly reduced. It could be seen that BMSCs transplantation could alleviate the infiltration of inflammatory cells and demyelination in the EAE mice.

### 3.6 Effect of BMSCs transplantation on inflammatory factor expression during EAE progression in mice

In the development of MS, TNF-α, IL-1β and IFN-γ played important roles in infiltration of inflammatory cells and demyelination, while IL-10 and TGF-β could alleviate the progression of MS. Therefore, we detected the expression levels of these inflammatory factors in the serum of mice in EAE group and EAE+BMSCs group by ELISA, and further detected their mRNA expression by RT-qPCR. ELISA results shown that after BMSCs transplantation, the levels of pro-inflammatory factors TNF-α, IL-1β and IFN-γ were significantly down-regulated, while the levels of anti-inflammatory factors IL-10 and TGF-β were reversed in serum of mice. In addition, RT-qPCR results shown that compared with the EAE group, the expression levels of pro-inflammatory factors TNF-α, IL-1β and IFN-γ in the EAE+BMSCs group were significantly down-regulated, while the expression level of anti-inflammatory factors IL-10 and TGF-β was significantly increased. These results were consistent with previous results of alleviating inflammation.

## Discussion

MS is a complex neuroinflammatory disease caused by local inflammation and immune dysfunction, leading to demyelination and extensive mononuclear cell infiltration[24–26]. Generally speaking, MS is considered as a disease of central nervous system involved in CD4^+^ T lymphocytes, including Th1 and Th17[27, 28]. Although the etiology still unclear, increasing evidence showed that genetic and environmental factors were associated with MS[29, 30]. It is well known that the complex pathogenesis of MS mainly includes demyelination and inflammation[31]. Transplantation or remyelination could promote the recovery of neurological function and demyelination of EAE[32, 33]. However, the transplantation of stem cells is the focus on the clinical treatment of MS[34, 35].

Increasing evidences have demonstrated that BMSCs promoted remyelinate axons and neurological function recovery in EAE animal model[36]. However, the mechanism of differentiation of BMSCs on EAE animal models remains to be studied. Myelination is carried out by oligodendrocytes in the central nervous system. Myelination is a modified expanded glial membrane that wraps around axons to achieve rapid saline-alkali nerve conduction and axon integrity[37]. Oligodendrocytes are derived from oligodendrocyte progenitor cells (OPC), and oligodendrocytes hold the capacity to proliferate, migrate and differentiate into myelinating oligodendrocytes. After demyelination, OPC is recruited to differentiate into myelinated oligodendrocytes, which then act on remyelination to protect axons from degeneration [38]. Therefore, oligodendrocytes are of great significance for remyelination. Astrocytes originate from neural embryonic progenitor cells arranged in the embryonic neural tube cavity[39]. It is currently recognized academically that astrocytes can support the function of oligodendrocytes. As early as 1984, studies have shown that type 1 astrocytes could expand O-2A progenitor cells from the optic nerve of newborn rats[40]. Later, Bhat and Pfeiffer observed[41] that extracts from astrocytes-rich cultures stimulated the differentiation of oligodendrocytes, thus supporting the concept that astrocytes played an active role in myelination. In addition, astrogliosis is one of the pathological features of MS. Astrocytes regulate the integrity of the blood brain barrier (BBB) by regulating the transport of peripheral immune cells[42], and can secrete a large number of chemokines and cytokines with pleiotropic functions[43], which lays an active role in promoting demyelination. So, this study investigated whether the transplanted BMSCs can differentiate into neurons, astrocytes and oligodendrocytes to affect the inflammatory invasion and demyelination of EAE mice. However, intravenous injection of BMSCs could not across the BBB, which is a key problem in the future. In the present study, BMSCs were injected into the lateral ventricle to detect the markers of oligodendrocytes, neurons and astrocytes. we found that BMSCs could be induced to differentiate into neurons, astrocytes and oligodendrocytes. Furthermore, BMSCs transplantation alleviated clinical signs, infiltration of inflammatory cells and demyelination of EAE mice, and down-regulated the expression levels of pro-inflammatory factors TNF-α, IL-1β and IFN-γ, while increasing the expression level of anti-inflammatory factors IL-10 and TGF-β.

Although stem cell transplantation is advanced, the success rate of stem cell transplantation is relatively low[44]. Some studies have evaluated the effect of stem cell transplantation on the clinical treatment of MS[45-47]. Thus, improving the success rate of stem cell transplantation could be used as a means of clinical treatment of MS.

## Conclusion

In conclusion, this study found that BMSCs could alleviate the inflammatory response and demyelination in EAE mice, which may be achieved by the differentiation of BMSCs into neurons, astrocytes and oligodendrocytes in EAE mice.

## Abbreviations

MS: Multiple sclerosis
EAE: Experimental autoimmune encephalomyelitis
BMSCs: Bone marrow mesenchymal stem Cells
CNS: Central nervous system
MAP-2: Microtubule-associated protein 2
GFAP: Gial fibrillary acidic protein
MBP: Myelin Basic Protein
O4: Oligodendrocyte Marker
LFB: Luxol Fast Blue

## Acknowledgements

We thank members of our laboratory for providing technical advice and encouragement.

## Authors’ contributions

Jia Geng conceived and supervised the study. Jia Geng and Guo-yi Liu a designed the experiment. Guo-yi Liu, Fan-yi Kong, Shu Ma and Li-yan Fu performed the experiments and analyzed data. Jia Geng wrote the manuscript with support from Guo-yi Liu and Fan-yi Kong. All authors read and approved the final manuscript.

## Availability of data and materials

The datasets used and/or analyzed during the current study are available from the corresponding author on reasonable

## Ethics approval and consent to participate

The animal experiments was approved by the Ethics Committee of Kunming Medical University and approved by Ethics Committee of Kunming Medical University

## Consent for publication

Not applicable

